# Uncovering immune characteristics of neurons in the context of multiple sclerosis and experimental autoimmune encephalomyelitis

**DOI:** 10.1101/2024.08.19.608118

**Authors:** Abdulshakour Mohammadnia, Sienna Sage Drake, Stephanie E.J. Zandee, David Gosselin, Jack P. Antel, Alexandre Prat, Alyson Fournier

**Affiliations:** Neuroimmunology Unit, Montreal Neurological Institute, Department of Neurology and Neurosurgery, McGill University, Montreal; Department of Neurology and Neurosurgery, Montréal Neurological Institute, McGill University, Montreal, QC, Canada; Department of Neuroscience, Faculty of Medicine, Université de Montréal, Montreal, QC, H3T 1J4, Canada; Department of Molecular Medicine, Faculty of Medicine, Université Laval, Quebec City, QC, Canada, G1V 4G2

**Keywords:** Experimental autoimmune encephalomyelitis, immune signature, multiple sclerosis, multi-omic data, neurons

## Abstract

Multiple sclerosis (MS) is an immune-mediated disease characterized by chronic inflammation and damage to the central nervous system, and substantial characterization of molecular signatures of glial and immune cells in the disease has been conducted. However, comparatively less well characterized is the molecular signature of pathologically inflamed neurons. Here, we accessed multi-omic high-throughput transcriptomic and epigenomic data to investigate the molecular signature of neurons from progressive MS patients and from mice subjected to experimental autoimmune encephalomyelitis (EAE). Our results indicate a shared and consistent neuronal gene expression signature in MS and EAE samples including a notable upregulation of immune system related genes and processes. Analysis of immune-enriched pathways in retinal ganglion cells (RGCs) and motor neurons from mice subjected to EAE revealed enrichment for the major histocompatibility complex class I pathway and interferon response genes in both types of neurons, and ATAC-seq analysis confirmed the accessibility of these genes. Parallel analyses of neurons from MS patients identified a signature of 48 commonly upregulated immune genes in neurons. Prediction of transcriptional regulators of these genes identified key upstream regulators including signal transducer and activation of transcription 1 (STAT1) and interferon response factor 5 (IRF5), highly involved in immune-related processes, with significant gene expression changes in both RGCs and motor neurons. Protein-protein interaction analyses among the transcriptional regulators unveiled intricate interactions among transcription factors, especially from the IRF and STAT families, suggesting their importance in regulating immune gene expression in inflamed neurons. Small molecule inhibition of IRF5 in inflamed neurons *in vitro* promoted cell survival and reduced the expression of the immune gene signature, supporting the biological relevance of IRF5 in mediating the neuronal inflammatory response. Together, this study identifies regulated immune processes in neurons from EAE mice and from MS patient samples suggesting potential roles for neuronal immune molecules in disease progression, as revealed by transcriptomic, epigenomic, and experimental approaches. Further, the results strongly implicate active interferon signaling in neurons driving immune related changes in gene expression through IRF5 and STAT1 transcription factor activity, potentially impacting downstream neuronal survival through IRF5 activity.

## Introduction

The hallmark pathological features of multiple sclerosis (MS) – central nervous system (CNS) inflammation, demyelination, and neurodegeneration – occur in tandem over the course of the disease, culminating in enduring clinical disability^1^. The cellular landscape mediating these changes in the CNS is remarkably diverse. Blood-brain barrier dysfunction facilitates the infiltration of peripheral immune cells and proinflammatory molecules from the bloodstream into the CNS parenchyma, creating a complex cellular milieu^2,3^. These peripheral immune components, including B-cells and T-cells drive inflammation and dysfunction of resident CNS cells encompassing astrocytes, microglia, oligodendroglial cells, ependymal cells, and neurons^4-6^. While extensive studies have explored the immune activity and signature of the immune system and glial cells in MS, the molecular signature of pathologically inflamed neurons remains a less-explored domain^7^.

Studies that have investigated neurons in MS have largely focused on describing axonal and neurite degeneration, neuronal cell dysfunction and death, and methods of promoting survival. Axonal and dendritic dysfunction has been described in EAE and MS, driven by membrane nano ruptures and ion channel dysregulation^8-10^. Wallerian degeneration likewise is involved in degeneration of chronically demyelinated axons through DLK1^11^. Other studies have demonstrated roles for autoantibodies and pathological protein accumulation, mitochondrial energy failure, altered potassium channel shuttling, and cellular senescence in inflamed neurons^12-16^. Additionally, in a single-nucleus sequencing study of MS patient brains there was a loss of *CUX2*+ cortical projection neurons identified^17^. More recently, work by Woo et al. (2024) characterized the role of stimulator of interferon response cGAMP interactor (STING) in orchestrating neuronal cell death during inflammation^18^. However, furthering our understanding of how neurons may integrate and respond to inflammatory signals in the context of MS is critical to the discovery of novel therapies that promote neuronal resilience to inflammation.

Here, numerous sequencing datasets across different affected neuron types including motor neurons (MNs) and retinal ganglion cells (RGCs) from murine experimental autoimmune encephalomyelitis (EAE) and cortical neurons from MS patients were leveraged to unravel gene expression and cellular pathway alterations occurring within neurons in response to injurious CNS inflammation. Somewhat surprisingly, a broad inflammatory gene expression signature was uncovered across these diverse neuronal datasets. Bioinformatic prediction of transcription factors upstream of this neuronal immune gene signature identified interferon response factor 5 (IRF5) and signal transducer and activation of transcription 1 (STAT1) as potential transcriptional regulators. Interestingly, small molecule inhibition of IRF5 promoted neuron survival in vitro. Together these data provide strong evidence for the interferon response pathway operating in neurons, regulating neuronal expression of immune genes, and invoke a role for the transcription factor IRF5 in regulating cell death in neurons.

## Results

### Immune Pathways Enriched in Neuronal EAE Models

The molecular signature of neurons exposed to pathological inflammation was studied in sequencing datasets of neurons from EAE mice and from MS patients. EAE data included bulk RNA-sequencing and ATAC-sequencing datasets from RGCs and a translating ribosome affinity purification RNA-sequencing dataset from MNs (**Fig.1A**)^15,16^. Single nuclei RNA (snRNA)-sequencing datasets from three independent human studies were analyzed in parallel to compare gene expression signatures in MS neurons and in the context of MS lesion pathology (**Fig.1B**)^17,19,20^.

**Figure 1.**
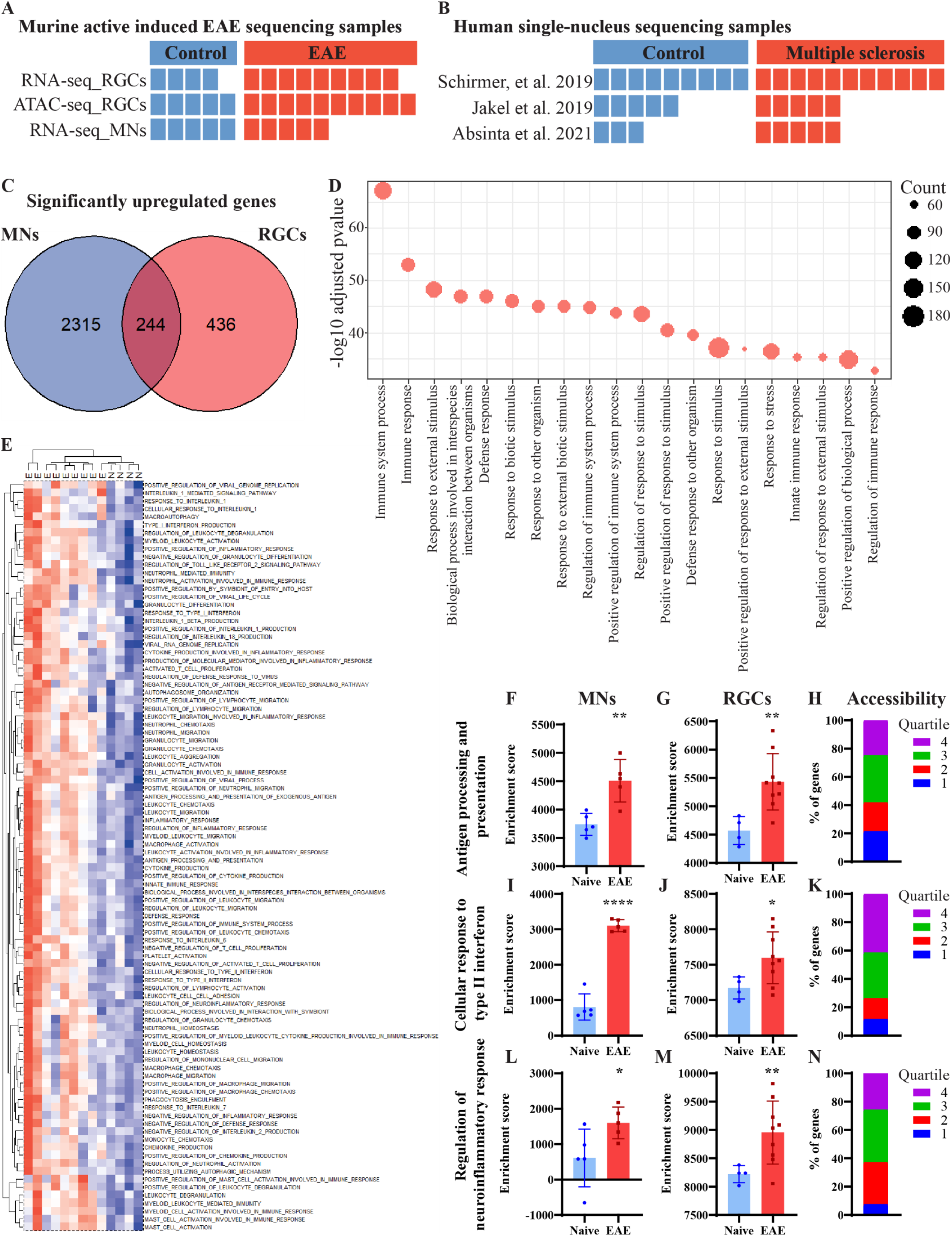
Immune processes significantly upregulated in retinal ganglion cells and motor neurons of EAE models. **A-B)** Details of the sequencing datasets incorporated into this study include multi-modal sequencing datasets of neurons from murine EAE **(A)** and single-nuclei RNA-sequencing datasets subset to neuronal populations from human MS patient and controls **(B)**. **C)** Venn diagram illustrates common upregulated genes in EAE samples versus control samples, based on p-value < 0.05 from each comparison (RGCs_EAE vs RGCs_Control, Motor neurons_EAE vs Motor neurons_Control). **D)** Gene ontology analysis of 244 common upregulated genes from EAE RGCs and MNs comparisons using g:Profiler web tool. Top biological processes were ranked based on adjusted p-values, and the top 20 pathways were visualized. Bubble sizes reflect the number of genes, and values on the y-axis represent the significance. **E)** Single-sample gene set enrichment analysis was applied to normalized read counts of EAE RGCs (labeled E) and naïve RGC samples (labeled N) to identify dysregulated immune processes. GO biological processes (GOBP) were filtered based on the enrichment scores higher than 5000 and p-values lower than 0.05. Red and blue colors indicate higher and lower enrichment scores, respectively. **F-N)** From ssGSEA analysis, examples of enriched GOBP-terms and their relative enrichment in MNs (**F,I,L)** or RGCs (**G,J,M)** and the percent of genes in each pathway belonging to which ATAC-sequencing quartile based on the normalized read count within 10k bp of the gene’s TSS (**H,K,N)**.

Analysis of RNA-sequencing datasets from murine motor neurons (MN) and retinal ganglion cells (RGC) identified numerous genes that were upregulated in both types of neurons in EAE compared to controls (*p-value* < 0.05, logFC > 0) (**Fig.1C**). To gain insight into the functional role of these genes, Gene Ontology (GO) analysis was conducted on the overlapping significantly upregulated genes between MNs and RGCs (**Fig.1D**). This analysis prioritized biological processes closely related with the immune system, such as “Immune system process”, “Immune response”, and “Innate immune response” (**Fig.1D)**. To produce a more detailed understanding of the immune signatures via GO analysis, single-sample gene set enrichment analysis (ssGSEA) was carried out on both datasets to assess the plurality of gene expression alterations at a pathway level. This revealed significant enrichment of an extensive spectrum of highly enriched immune pathways in EAE RGCs compared to their naïve counterparts (p-values < 0.05, enrichment score > 5000) (**Fig.1E**). This included enrichment of pathways related to viral defense and interferon signaling including the GO terms “cellular response to type II interferon”, “type I interferon production”, and “response to type I interferon”. Similar trends were noted in EAE MNs in comparison with naive MNs (**Fig.S1**). Indeed, there were multiple pathways significantly enriched in both EAE RGCs and MNs such as “antigen processing and presentation”, “cellular response to type II interferon signaling”, and “regulation of neuroinflammatory response” (**Fig.1F-N).** Using RGC ATAC-sequencing data, all genes in the genome were categorized into quartiles based on the normalized read counts within 1 kbp of their transcription start site (TSS), and the significantly regulated immune pathways were assessed based on the accessibility of their member genes, ranging from least accessible (quartile 1) to most accessible (quartile 4). Here, a majority of genes in these immune pathways belonged to the top two quartiles indicating a high level of accessibility (**Fig.1H,K,N, Fig.S2)**. Together, these data demonstrate that neurons from mice subjected to EAE express a complex transcriptomic profile of immune genes that are accessible and common to both MNs and RGCs.

### Immune Processes Enriched in Human Multiple Sclerosis Samples

Expanding the investigation to human MS samples, a dataset comprised of snRNA-sequencing of gray matter from MS patients and controls was reanalyzed (**Fig.1B)**^17^. This analysis including clustering and annotating cell identities, to identify excitatory and inhibitory neurons in MS patients and controls (**Fig. S3A-D)**. Gene expression data was normalized and averaged by patient sample for all neuron assigned nuclei, and ssGSEA was conducted for the immune pathways identified as significantly enriched in EAE RGCs. Of these pathways, eleven were likewise significantly enriched (p-values < 0.05) in MS versus control patient neurons **(Fig.2A)**. Indeed, among those pathways were again “type I interferon production” and “antigen processing and presentation”. The genes annotated to these pathways were compared to the significantly upregulated genes from EAE MNs and EAE RGCs, and a signature of 97 genes common to the human MS neuron pathways, and the two EAE neuron datasets was identified **(Fig.2B)**. Individual genes from this list of 97 were assessed for ATAC-seq accessibility, which showed unequivocal evidence of peaks at the TSS of *H2-D1*, *Calr*, *B2m*, *Stat1*, and *Psmb8* (**Fig.2C)**. Of these five genes, four are significantly upregulated in human MS patient neurons compared to controls – *B2M*, *CALR*, *HLA-C*, and *PSMB8* (**Fig.2D-G)**. These same genes had high and increasing expression in the EAE MNs and RGCs (**Fig.2H-O)**. Leveraging another two snRNA-sequencing datasets from MS patients, signatures from nuclei classified as neurons were compared across different MS lesion types. Here, assessment of the 97 gene signature found robust enrichment of many of these genes in neurons from chronic active lesions, suggesting that ongoing local inflammatory activity may be driving upregulation of these genes in neurons (**Fig.2P,Q)**. Altogether, these data demonstrate conserved enrichment of immune pathways and genes in diverse neuron types in EAE and in cortical neurons from MS patients, associated with inflammatory lesion activity.

**Figure 2.**
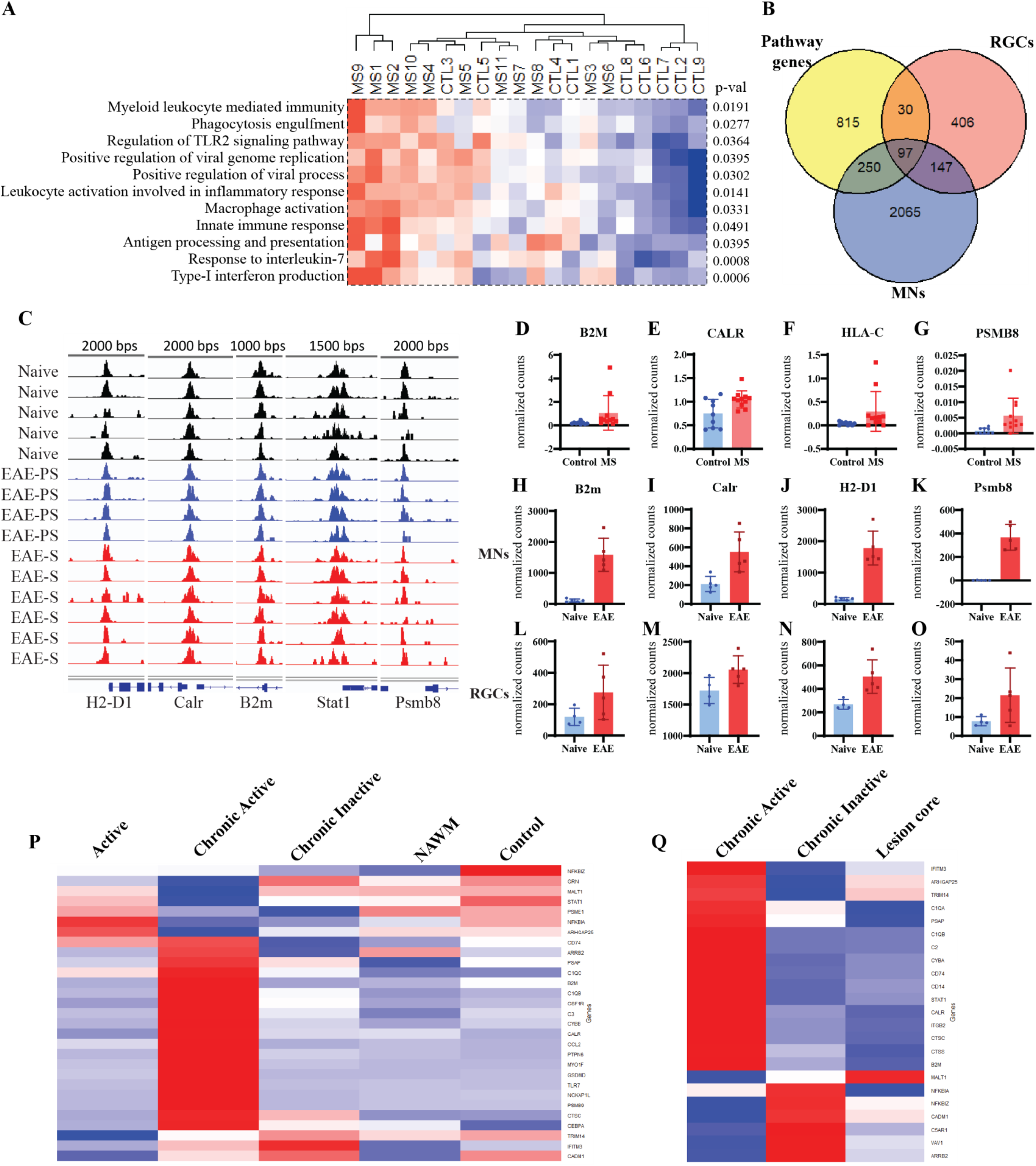
A common immune gene signature between MS patient neurons and murine EAE neurons. **A)** Pathway enrichment analysis of cortical neurons from control and MS patients reveals significant enrichment of immune related pathways that are also significantly enriched in RGCs from EAE mice. **B)** Venn diagram showing the overlap between the genes belonging to the 11 significantly enriched pathways in human MS neurons, and significantly upregulated (p <0.05) genes from EAE RGCs and MNs. Red and blue colors indicate higher and lower enrichment scores, respectively **C)** ATAC-seq genomic tracks demonstrating high accessibility at transcription start sites across naïve, presymptomatic EAE (EAE-PS) and symptomatic EAE (EAE-S) ATAC-sequencing samples for genes from the 97 gene signature. **D-G)** Normalized read counts from neurons from single-nucleus sequencing dataset of MS patients and controls for notable genes from the 97 gene signature demonstrating a trend towards upregulation. **H-O)** Normalized read counts from MNs (**H-K)** and RGCs (**L-O)** of the same genes from the 97 gene signature demonstrating upregulation across these neurons. **P-Q)** Analysis of neurons from another single-nucleus datasets of MS patients (**P:** Absinta et al. 2021,**Q:** Jakel et al, 2019) where samples are classified based on lesion type demonstrates a strong enrichment for members of the 97 gene signature in chronic active lesions. Red and blue indicate higher and lower expression levels, respectively.

### Prediction of upstream regulatory transcription factors governing immune gene expression in neurons

The 97 gene signature was subset based on ATAC-seq accessibility, and the top accessible genes (n=48) were visualized in a network, demonstrating a high level of interconnectivity between these genes (**Fig.3A)**. Next, using iRegulon, a list of transcription factors predicted to regulate the genes was generated and incorporated into a regulatory network for centrality analysis. Here, the top regulated genes were visualized as those with the highest level of in-degree connections (**Fig. 3B)**. Many of the prioritized genes were previously described as highly regulated and accessible, such as *B2m* and *Stat1* (**Fig.3B)**. Similarly, the top regulators were visualized based on their high level of out-degree connections, prioritizing numerous transcription factors belonging to the nuclear factor kappa beta (NFκβ) and IRF families, all regulating around 30 genes of the 48 gene signature (**Fig.3C)**. To better understand transcription factor cooperation in regulating the immune gene signature, all predicted upstream regulators were loaded into StringDB to identify protein-protein interactions between these factors. This network was subsequently analysed by Leiden clustering in Cytoscape to identify highly cooperative transcription factor subnetworks. Here, a large, interconnected subnetwork was identified, with notable contribution from the IRF and STAT family transcription factors; another subnetwork of interest contained many members of NFκβ family (**Fig.3D, FigS4)**. Importantly, many of the transcription factors in the primary subnetwork were significantly upregulated in one or both EAE neuron RNA-sequencing datasets, including *Stat1*, *Stat6*, *Spi1*, and *Irf5* (**Fig.3D)**. Next, a list of genes associated with MS susceptibility was compared to the list of predicted upstream regulators. A single transcription factor belonged to both sets, IRF5, which is predicted to regulate numerous members of the 48 gene signature (**Fig.3E,F)**. Together, these data identify a cohesive set of 48 genes associated with neuronal inflammation, high confidence transcription factor regulators of this signature, and an emphasis of IRF and STAT transcription factors as key players in their regulation.

**Figure 3.**
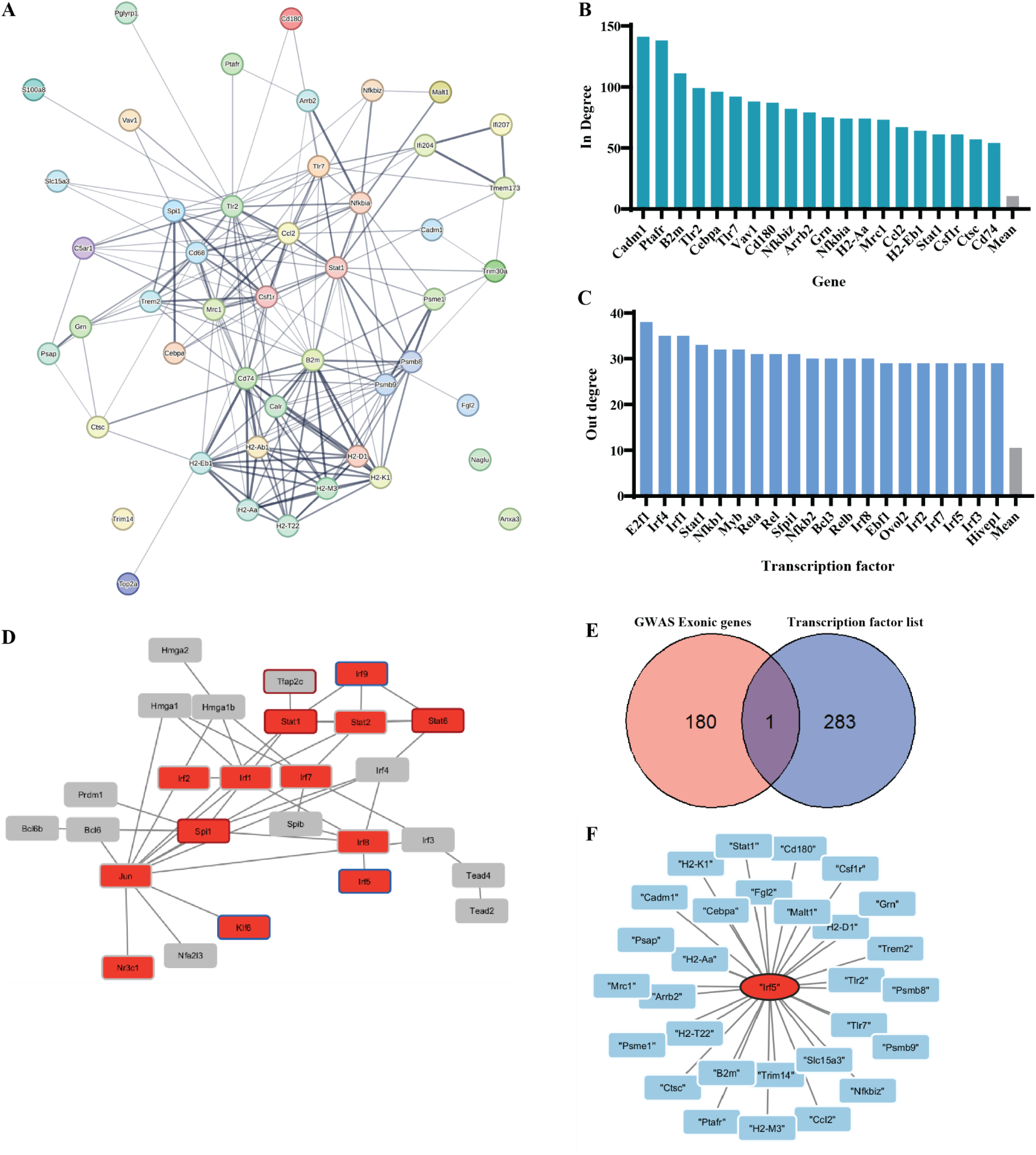
Transcription factor regulatory network of prioritized neuronal immune genes. **A)** Visualization of 48 genes used for upstream prediction of transcription factors using iRegulon. These genes were selected from the 97 genes common to human MS neuron enriched pathways and upregulated in EAE MNs and RGCs and narrowed down based on evidence for accessibility in RGCs from ATAC-seq data. **B,C)** Centrality analysis of the predicted transcription factor – 48 genes interaction network using CentiScape tool in Cytoscape. Visualization of in degree (**B)** demonstrates the top 20 regulated genes, while out degree (**C)** demonstrates the top 20 regulatory transcription factors. **F)** Protein-protein interaction analysis of transcription factors using String to identify PPIs and Leiden clustering algorithm to prioritize most interconnected sub-network. Red fill indicates significant upregulation in EAE MN samples compared to control, while a red border indicates significant upregulation in EAE RGCs. A blue border indicates a near significant upregulation in RGCs (p < 0.06). **E)** Comparison of MS susceptibility genes identified in a large GWAS cohort with transcription factors predicted to regulate immune genes in EAE models reveals a single transcription factor common to both groups. **F)** Visualization of the interaction network for the single transcription factor from E: Irf5. Blue nodes are genes from the initial list of 48 that are predicted to be regulated by Irf5.

### IRF5 inhibition protects neurons from inflammation mediated cell death *in vitro*

To investigate whether this bioinformatic approach combining numerous different types of sequencing data could indeed coalesce on important regulators of neuronal inflammation, an *in vitro* approach was used to further investigate IRF5 due to its upregulation in EAE, relevance to MS susceptibility, and high number of predicted regulated immune genes. Here, primary mouse cortical neurons were cultured for 7 days before being exposed for 72h to condition media from peripheral blood mononuclear cells stimulated with PMA-ionomycin (PBMC-CM). Cortical neurons received either control or PBMC-CM, and DMSO or 1 µM of the Irf5 inhibitor YE6144 supplemented in the media (**Fig.4A)**. PBMC-CM treatment induced significant cell death in 72h (**Fig.4B,C)**. However, cells receiving supplementation of 1 uM YE6144 demonstrated significantly higher neuronal survival in the PBMC-CM condition, without altering survival in the control media condition (**Fig 4.C)**.

**Figure 4.**
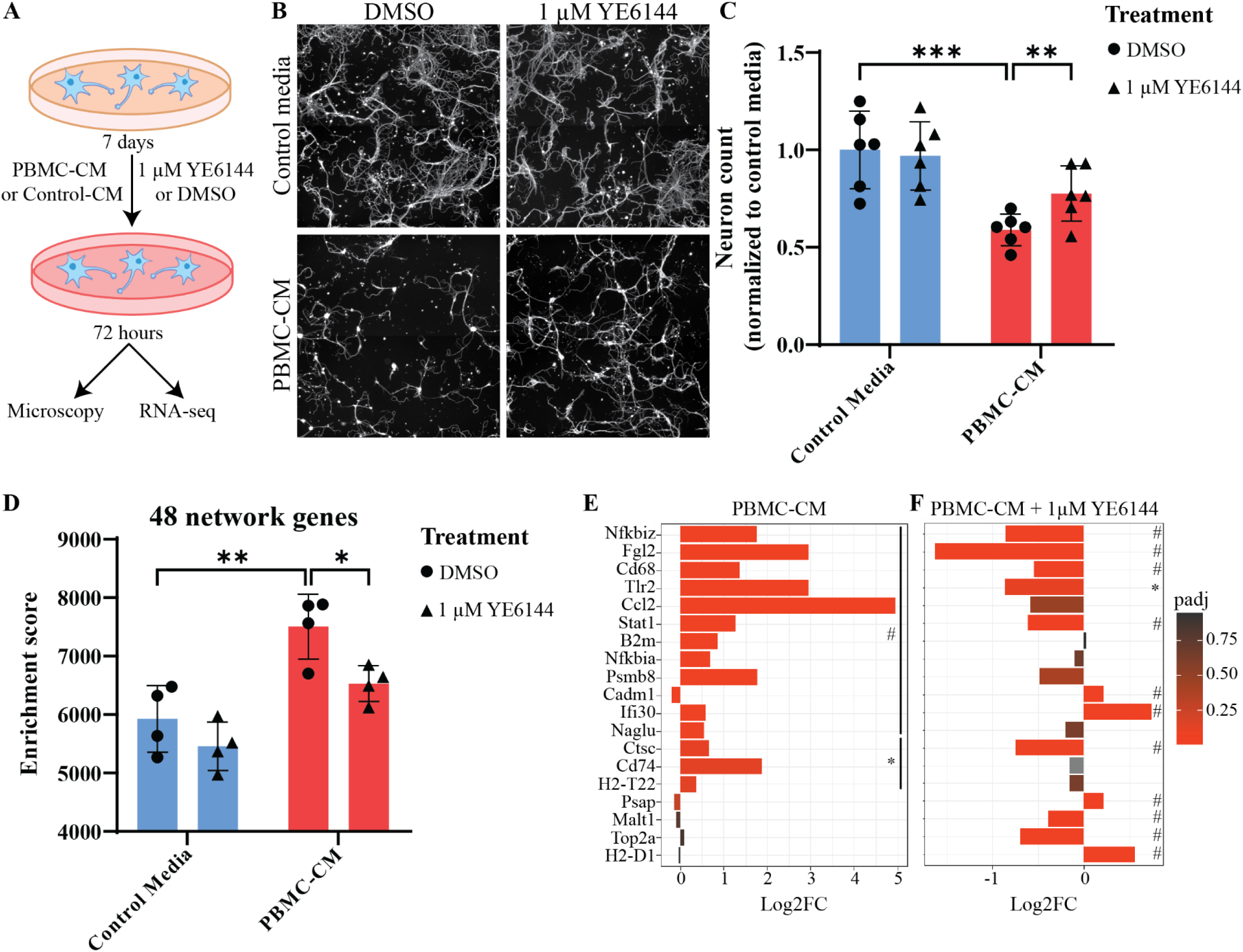
Small molecule inhibition of IRF5 protects neurons from immune-cell condition media induced cell death. **A)** Experimental design for testing the IRF5 inhibitor YE6144 on neuronal survival and gene expression *in vitro*. **B)** Representative images of cortical neurons cultured in the presence of control media or PBMC-CM, and DMSO or YE6144. **C)** Neuronal cell counts after treatment with control media or PBMC-CM, and DMSO or YE6144. ** p < 0.01, *** p < 0.001, Two-way Anova with Holm-Sidak multiple comparison correction. **D)** Single sample gene set enrichment score for the list of 48 genes from Fig. 3A from RNA-sequencing results of neurons after treatment with control or PBMC-CM and DMSO or YE6144. Two-way Anova with Holm-Sidak multiple comparison correction. **E,F)** Visualization of a subset of the 48 genes that are significantly dysregulated when comparing either PBMC-CM + DMSO to control media + DMSO, or PBMC-CM + YE6144 vs PBMC-CM + DMSO # = padj < 0.1, * = pval < 0.05.

Next, RNA-sequencing of primary mouse cortical neurons treated with and without PBMC-CM and YE6144 was conducted to assess the gene expression profile in this experimental paradigm. To understand whether PBMC-CM induced expression of immune genes similarly to the EAE and MS sequencing datasets, ssGSEA of the 48 network genes identified from the EAE and MS datasets was performed and found that the set of 48 network genes was enriched in the transcriptome of neurons that received PBMC-CM treatment. However, treatment with the IRF5 inhibitor YE6144 significantly reduced the enrichment of the 48 gene signature (**Fig. 4D)**. Additionally, assessment of individual genes from the network found a number of these were significantly upregulated with PBMC-CM treatment including *Stat1*, *Nfkbiz, Nfkbia, Fgl2, and Tlr2* (**Fig. 4E**). Similarly, many of these were significantly downregulated with YE6144 treatment, including *Stat1, Nfkbiz,* and *Fgl2* (**Fig. 4F).** All together, these data provide further validation that IRF5 regulates expression of inflammatory genes in neurons, and inhibition of IRF5 protects neurons from inflammation mediated cell death.

## Discussion

The data presented in this study provide evidence for upregulation of an inflammatory gene signature among neurons in EAE and MS. Particularly notable is the enrichment of genes and processes related to interferon signaling and antigen presentation. These features were accessible based on neuronal ATAC-sequencing data, and transcription factor prediction of a subset of commonly regulated immune genes prioritized upstream regulatory factors that mediate interferon signaling including STAT1 and IRF5. While interferon signaling activity has been previously described in glial cells in the CNS in MS, these data represent a novel characterization of a neuron’s role in potentiating inflammation in the disease^21-23^.

Broadly, the role of interferon signaling is to establish an antiviral state in cells in response to infection^24^. Outside of infection however, interferons exert a broad range of effects in neurons. Within the type 1 interferons, interferon-α has a nefarious effect, inducing dendrite retraction and cell death, while interferon-β is essential to neuronal growth and survival^25-27^. Given that both signal through the same receptors, IFNAR1 and IFNAR2, this suggests that the molecules have divergent outcomes for downstream effectors in neurons. Meanwhile, type 2 interferon-γ (IFNγ) has complex roles in the neuronal antiviral response; inducing neuronal viral clearance while bypassing cell death^28^. In non-infected conditions, loss of IFNγ promotes hippocampal and synaptic plasticity in learning, which pairs well with the fact that IFNγ application in cultured neurons inhibits neurite outgrowth and synaptic development and facilitates dendrite retraction^29,30^. IFNγ signals through its cognate receptor complex of IFNGR1 and IFNGR2 triggering STAT1 phosphorylation, transcription factor complex formation with IRFs, and expression of interferon response genes (IRGs) including antigen presentation genes^24,31-33^. Intracellular signaling elements, including STING mediated cytosolic DNA sensing and toll-like receptors can likewise induce STAT phosphorylation and IRF activity resulting in increased IRG expression^24,34^. This is particularly relevant in neurons during inflammation where STING orchestrates release of intracellular calcium stores resulting in neuronal dysfunction and death in response to stressors including IFNγ and glutamate^18^.

Moreover, while neurons do not typically express major histocompatibility complex (MHC) molecules in the healthy adult brain, stimulation *in vitro* with IFNγ has been shown to induce MHC molecule expression on neurons. Similarly, suppression of electrical activity in neurons further promotes MHC expression *in vitro*, suggesting a possibility that interference with the homeostatic electrical activity of neurons may induce the antigen presentation pathway^35^. Multiple studies of post-mortem tissue in aging and disease – particularly in virus infection – find the presence of MHC molecules on neurons^36^. Intriguingly, studies have likewise reported elevated MHC-I antigen presentation protein expression on neurons in MS lesions and expression of the MHC-I gene regulatory transcription factor NFκβ p65^37,38^. Interferon signaling and antigen presentation are intimately associated, and MHC molecules on neurons has been shown to lead directly to T-cell mediated neurite injury, thus these pathways are likely to be critical components of the neuronal response to inflammation in the context of MS^39,40^.

Meanwhile, the role of IRF5 remains obscure, although bioinformatic prediction positions it as a top regulatory element of the immune genes activated in inflamed neurons, and GWAS studies have found a genetic association with it and MS susceptibility^41^. The very interesting result that its inhibition promoted neuronal survival in the context of *in vitro* inflammatory insult suggests that perhaps it promotes activation of cell death pathways downstream of inflammatory insults. IRF5 has known roles in systemic autoimmunity, but its expression and role in neurons remains understudied^42,43^. Only one other study has assessed a role for IRF5 in neurons; here too, IRF5 inhibition was protective in a motor neuron like cell line overexpressing a truncated form of TDP-43^44^. Indeed, IRF5 is known to be essential to apoptosis in other cell types, and its knockout results in resistance to apoptosis in hepatocytes and dendritic cells^45^.

Future studies into neuronal immune signaling in MS and its models would benefit from more detailed investigation of individual molecules and receptors in these pathways to delineate specific modulatory effects this complex network of associated genes can have in inflammatory injury. Additionally, while the data presented here benefited from the investigation of multiple different neuronal subtypes to identify a common immune signature, it is highly likely that varied neuronal subtypes have varied responses to inflammation. Perhaps susceptible subsets have higher baseline expression of IRF5 or other pro-apoptotic factors, while more resilient subtypes utilize different signaling mechanisms allowing them to evade cell death. Thus, this remains an important area for future investigation.

### Funding and acknowledgments

S.S.D. received funding from the Canadian Institutes for Health Research (CIHR) Vanier Canada Graduate Scholarship. A.E.F. is funded by the CIHR and MS Canada. We would like to acknowledge the support of the McGill Advanced Bioimaging Facility for their expertise in microscopy and access to the high-content screening microscope.

## Materials and methods

### ATAC-sequencing data processing

Raw reads were trimmed with trimmomatic, aligned to the genome with bowtie2, and quantified within 1,000 bp of annotated transcription start sites in VisR to obtain normalized read counts across genes as previously described^16,46^. The normalized counts across genes were then grouped into quartiles from lowest reads (Quartile 1) to highest reads (Quartile 4). Individual tracks were additionally loaded into the UCSC browser for visualization to identify bona fide TSS peaks.

### RNA-sequencing data processing

Raw RNA-sequencing data was accessed through the GEO^15,16^. Reads were trimmed with cutadapt, aligned with hisat2, and count matrices were generated by htseq. DESeq2 was used for count normalization and differential gene expression analysis^47^. GProfiler was used to determine enriched pathways based on significantly upregulated genes^48^. Single sample gene set analysis conducted with GSVA package on the normalized counts table for each dataset^49^.

### Single-cell sequencing data analysis

The single-nucleus RNA sequencing (snRNA-seq) data in this study were obtained from a previously published dataset by Schirmer et al. in 2019, accessible under accession number PRJNA544731 (NCBI Bioproject ID: 544731) and processed as previously described^16,17^. Cell types were annotated using reference publication of Schirmer et al., datasets and clusters renamed based on their cell identities. Then, the normalized average gene expression was compiled across neuron clusters and stratified by patient. Pathway analysis was conducted with GSVA package for single-sample GSEA on the normalized counts table. Additional single-nucleus sequencing datasets (GSE118257, GSE180759) were accessed and processed in the same manner, then annotated further based on lesion classification in the sample metadata^19,20^.

### Network visualization and transcription factor prediction

Network visualizations were made in String and Cytoscape. Transcription factor prediction was conducted on the list of 48 genes using iRegulon with standard settings. The Centiscape plugin for Cytoscape was used on the entire network of transcription factors and their target genes for centrality analysis. The list of identified transcription factors was visualized in String to obtain protein-protein physical interactions, then loaded into Cytoscape for Leiden clustering for subnetwork identification. GWAS associated genes were filtered on exonic associations only^41^.

### Peripheral blood mononuclear cell condition media preparation

Blood from adult Sprague–Dawley rats was collected by cardiac puncture as previously described following an animal use protocol approved by the Montreal Neurological Institute animal care committee according to Canadian Council on Animal Care guidelines^50^. PBMCs were separated by Ficoll-Paque density centrifugation followed by 3 PBS washes. After the second wash, cells were incubated for five minutes in red blood cell lysis buffer, then washed for the third time. Cells were plated in 6 well plates containing Neurobasal media with 1% glutamine/1% penicillin/streptomycin. PBMCs were either left unstimulated or activated for 3 days by adding phorbol 12-myristate 13-acetate (PMA) and ionomycin (Sigma-Aldrich Canada, Oakville, ON) (P/I) at final concentrations of 0.08 μg/mL PMA and 1 μg/mL ionomycin. Control media was cultured for the same time and with the same PMA/I supplementation, but without cells.

### Mouse cortical neuron culture and immunocytochemistry

All animal experiments were performed according to the guidelines set by the Canadian Council on Animal Care (CCAC). Cortical neurons were isolated and cultured as previously described^51^. Briefly, E16 pregnant females were sacrificed, and embryos dissected to isolate cortices. These were digested for 30 minutes at 37°C with trypsin, then washed, and plated on PLL coated 96 well plates in DMEM/FBS. After 1 hr, the media was swapped to Neurobasal with 1% N2/ 1% B27. Cells were maintained in Neurobasal for 7 days, at which point a half media swap was conducted with either control media or stimulated PBMC-CM, containing DMSO or a final concentration of 1 uM of YE6144 in DMSO (MedChem Express, Cat. No.: HY-150095). Neurons were left for 3 days, then fixed with 4% PFA/20% sucrose in PBS for 30 minutes at room temperature. Following fixation, cells were washed three times with PBS, blocked and permeabilized with 0.2% Triton-100/5% BSA in PBS for 1 hr at room temperature, then incubated in primary antibody against betaIII-tubulin (Cell Signaling) for 1 hr at room temperature in 0.2% Triton/1% BSA. Following primary antibody incubation, cells were washed 3 times with PBS, and incubated for 1 hr in secondary antibody (Thermofisher) and Hoechst dye at 37°C, before a final set of washes with PBS. Images were acquired on a high content screening microscope, using a 10X objective and 5 ROIs per well. Image quantification was conducted in ImageJ, using a custom ImageJ macro to count the number of beta-III-tubulin positive nuclei.

---

**Fig S1.**
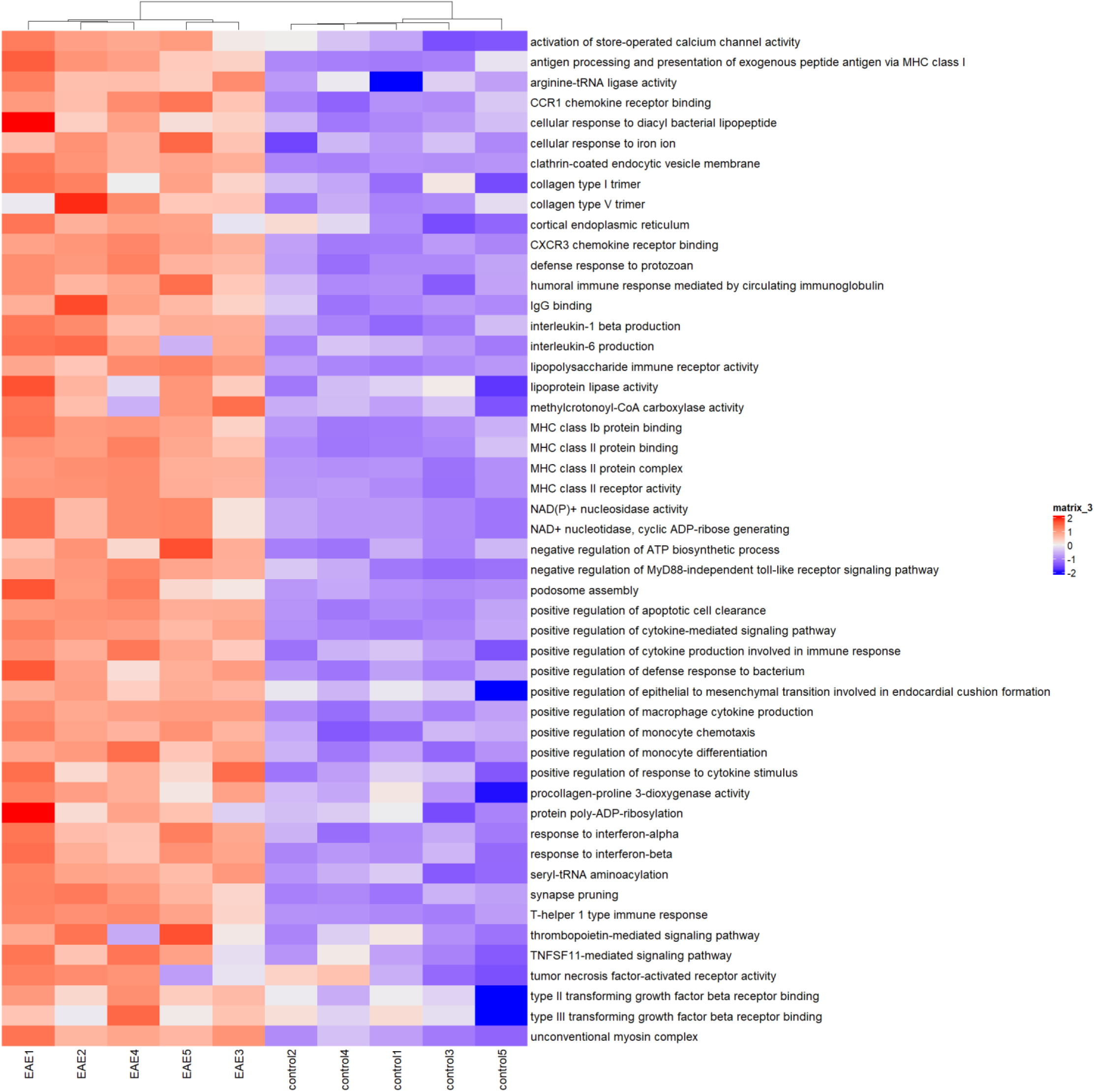
Motor neuron top 50 enriched pathways.

**Fig S2.**
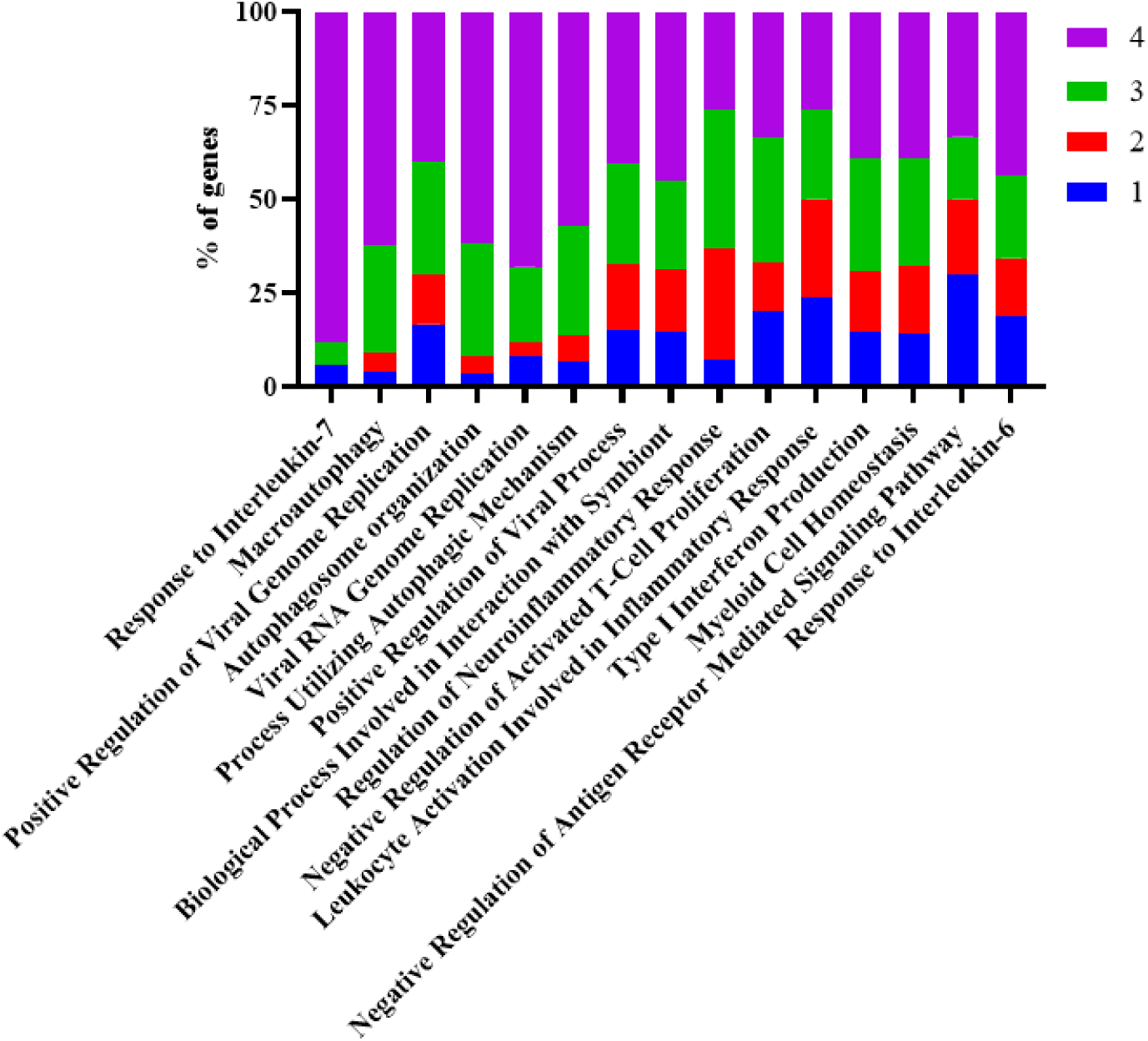
Accessibility of fifteen top RGC pathways.

**Fig. S3.**
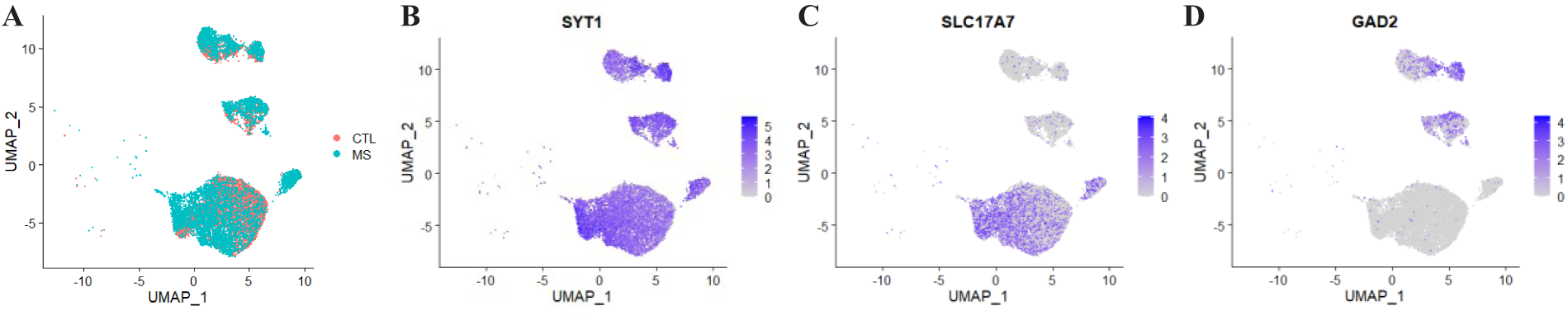
UMAP representation of single cell data from Schirmer et al. 2019.

**Fig. S4.**
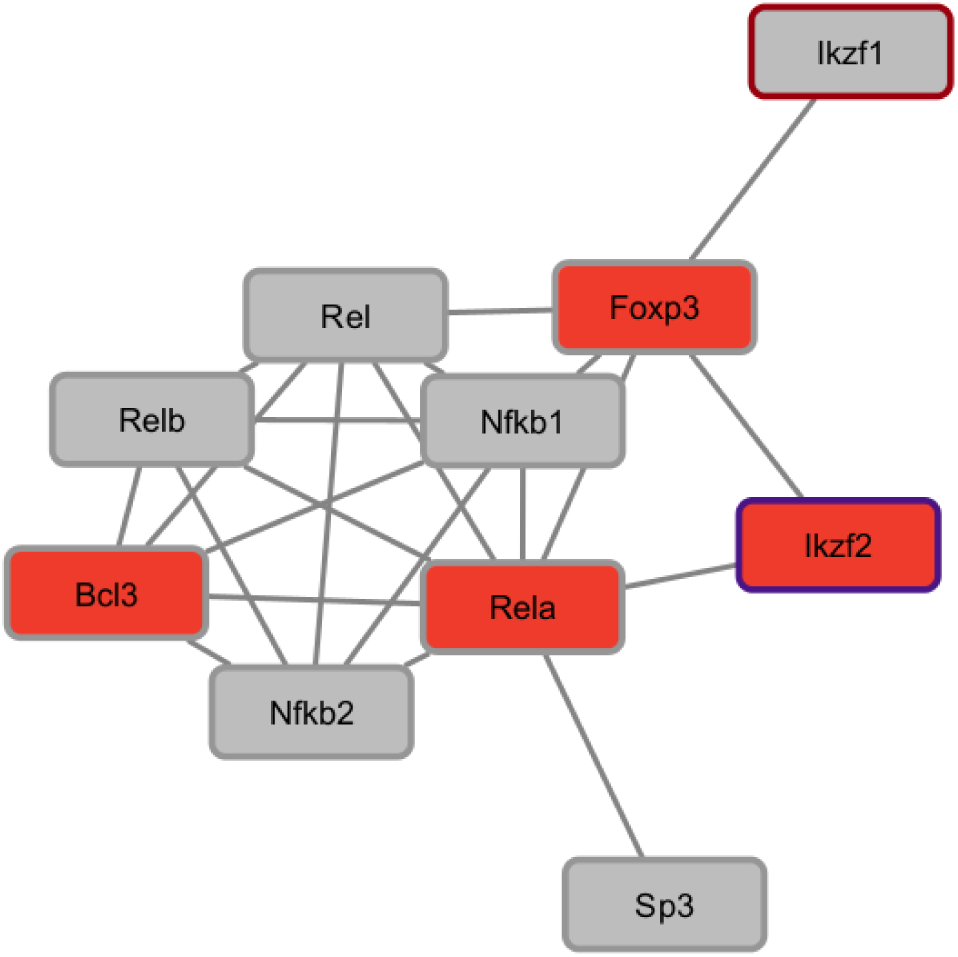
Second largest transcription factor subnetwork. Red fill indicates significant upregulation in EAE MNs, red border indicates significant upregulation in EAE RGCs, purple border on Ikzf2 indicates its significant downregulation in EAE RGCs.

